# SNAzzy: an image processing pipeline for investigating global Synchronous Network Activity

**DOI:** 10.64898/2026.11.05.764570

**Authors:** Carlos D. Paiva, Alana J. Evora, Shirui Zheng, Arnaldo Carreira-Rosario

**Affiliations:** Department of Biology, Duke University, Durham, NC 27708; Department of Neurobiology, Duke University, Durham, NC 27708

## Abstract

Genetically encoded fluorescent indicators are powerful tools for monitoring biological processes in live samples (Lin and Schnitzer 2016; Nakai, Ohkura, and Imoto 2001). When combined with a large field of view, a single time-lapse recording has the potential to capture many specimens, facilitating high-throughput data collection. However, this approach generates large, multidimensional datasets that are challenging to process and analyze. We present `SNAzzy`, a Python package for studying synchronous network activity (SNA) in Drosophila embryos via high-throughput microscopy. SNA is a hallmark of developing nervous systems (Wu, Kourdougli, and Portera-Cailliau 2024; Blankenship and Feller 2009; Akin and Zipursky 2020), often studied using genetically encoded calcium indicators to monitor neural activity in vivo. `SNAzzy` processes and analyzes time-lapse datasets taken from live samples using fluorescent widefield microscopy. Each dataset contains dozens of individual specimens in the same field of view and thousands of time points. The software offers individual specimen cropping for optimization of storage and processing, adaptive regions of interest for quantification of fluorescence and changes in morphology over time, a custom peak detection algorithm, and a graphical user interface for data visualization, curation, and dataset comparison. The code is available at https://github.com/ACRLab/snazzy.

## Statement of Need and State of the Field

During synchronous network activity (SNA), many neurons fire simultaneously, generating waves of activity that span large portions of the nervous system (Blankenship and Feller 2009; Wu, Kourdougli, and Portera-Cailliau 2024; Akin and Zipursky 2020). In Drosophila embryos, SNA typically lasts 4 hours, during which the nervous system undergoes a stereotyped morphological change via ventral nerve cord condensation (Crisp et al. 2008; Carreira-Rosario et al. 2021; Karkali et al. 2022). To gain an understanding of SNA, it is essential to quantify waves of activity in the nervous system while also tracking morphology as a proxy for neurodevelopment. For these reasons, we combine a commonly used genetically encoded calcium indicator (GECI) that reports neural activity with a structural fluorophore (Carreira-Rosario et al. 2021). The structural fluorophore signal remains stable, independent of neural activity, making it suitable for continuously tracking morphology of the nerve cord. To record many embryos during SNA, we use a wide-field fluorescence microscopy system that captures the GECI and structural fluorophore signal of dozens of developing embryos for over 5 hours.

We were unable to find a tool designed for widefield microscopy that rapidly processes multiple specimens, quantifies levels of fluorophore activity, and incorporates a peak-finding algorithm suitable for global calcium traces. `SNAzzy` is designed to investigate global levels of neural activity across multiple developing embryos simultaneously.

### Tracking of multiple “adaptive ROIs”

To the best of our knowledge, there are no other packages that provide functionality for automated parsing of raw images of many live specimens into activity and morphological quantifications. Other studies have employed manual selection of regions of interest (ROIs) and used static ROIs (Akin and Zipursky 2020; Menzies et al. 2024; Ardiel et al. 2022; Carreira-Rosario et al. 2021). Manual selection often generates imprecise ROIs, which can lead to inaccurate quantifications, and is also cumbersome and prone to human error. Static ROIs are not reliable for detecting the fluorescent signal of live specimens that change in morphology and move while imaging. `SNAzzy` fills these gaps as an accessible pipeline for the automated analysis of multiple live samples in parallel. The pipeline generates an “adaptive ROI” that changes frame-by-frame for each specimen. This enables the accurate tracking of fluorescence intensity as well as changes in tissue morphology or size. `SNAzzy`’s design provides an automated, modular, and fully auditable workflow, and ultimately contributes to more reproducible and comparable results across experiments.

### Capturing global calcium dynamics

To the best of our knowledge, there are no open-source packages that provide tools for performing automated data analysis and quantification of global calcium dynamics. Most open-source tools available for analyzing neural activity using GECI focus on segmenting individual neurons within a single specimen. `CaImAn` (Giovannucci et al. 2019) and `Suite2p` (Pachitariu et al. 2016) are among the most widely used. These packages detect calcium dynamics and use individual neuron statistics to perform spike inference, but do not offer direct peak detection on the calcium signal. Furthermore, they are optimized for two-photon microscopy as opposed to wide-field microscopy. `SNAzzy` provides a series of automated analyses and quantifications to analyze global calcium levels in time-series acquired with widefield microscopes.

## Research Impact Statement

SNAzzy is a complete solution for automated analysis of calcium dynamics from high-throughput time-lapse microscopy datasets. SNAzzy has had a significant impact on our laboratory by speeding up the analysis and onboarding researchers with little to no prior Python experience. This software replaces the previous, manually intensive analysis pipeline (Carreira-Rosario et al. 2021) with a fully automated workflow. Compared to the manual workflow, SNAzzy offers additional tools and reduces analysis time from days to minutes. Several students with limited programming background have installed SNAzzy and analyzed hundreds of specimens within a week. Lastly, lab members have added modules to SNAzzy to address important biological questions. This has led to new biological insights and the development of ongoing research projects.

Given that our lab is pioneering the study of SNA in Drosophila embryos, and that SNAzzy is designed as a generalizable and extensible analysis framework, we expect near-term adoption by other laboratories studying SNA and related developmental imaging paradigms. Lastly, beyond Drosophila embryos, the code can be used to support automated quantification of global fluorescence dynamics in other experiments that involve simultaneous imaging of multiple samples, particularly small-sized specimens (Donoughe, Kim, and Extavour 2018; Avasthi et al. 2023; Yamamoto and Yuste 2023).

## Software Design

`SNAzzy` consists of two packages: a processing and an analysis package. The former contains an image processing pipeline, while the latter produces quantifications and visualizations. This separation of concerns promotes modularity and extensibility, making it easy for users to repurpose the packages elsewhere.

The processing package implements an image processing pipeline composed of modular stages. The pipeline converts large image files into lightweight CSV files containing fluorescence intensity traces and shape metrics, thereby substantially reducing storage requirements and facilitating data sharing and reuse. The analysis package consists of a core Python library and an interactive Graphical User Interface (GUI) implemented using `PyQt6`. The core library performs peak detection, metric extraction, and statistical analyses, while the GUI provides tools for visualization, curation, and comparison across datasets. Communication between the GUI and the core analysis code is mediated by a Model layer, which stores all data required for presentation and keeps the core code lean.

Importantly, all experiment-specific parameters, including peak detection and signal-processing settings, are stored in a single JSON configuration file. This design ensures that analyses are fully reproducible and that configurations can be easily shared across users, machines, and experimental replicates.

Overall, our software design decisions focus on supporting reproducible research practices and making it easy to extend the pipeline to new imaging modalities, metrics, or biological systems.

## Pipeline Description

The initial input for `SNAzzy` (Figure 1) is raw time-lapse imaging data containing multiple embryos. Each embryo expresses a GECI (dynamic fluorophore) and a structural fluorophore. Fluorophores are imaged in different optical channels.

**Figure 1:**
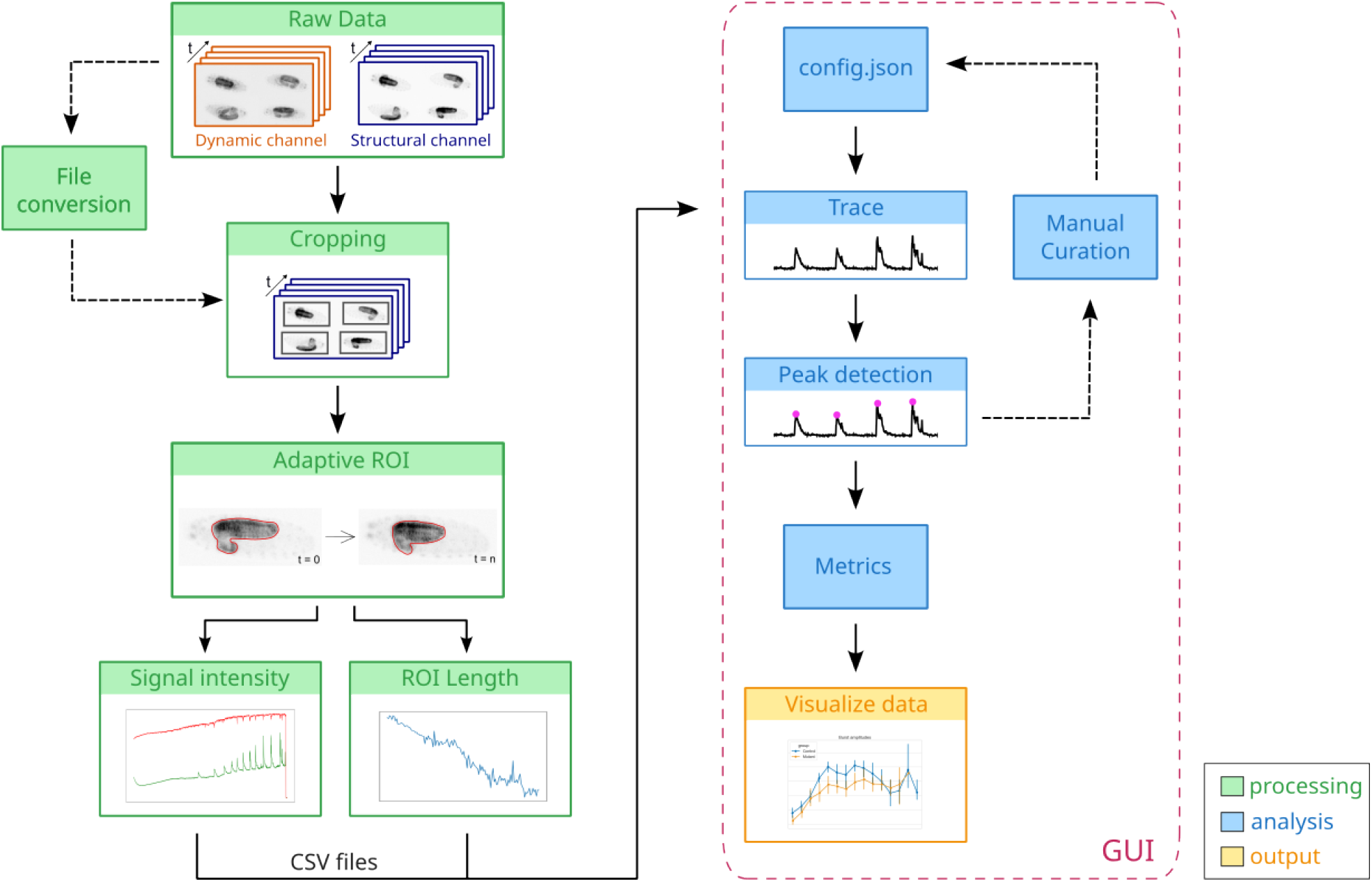
Schematic of the SNAzzy pipeline. Time-lapse recordings from fluorescent widefield micro-scopes (raw data) enter the processing stage (green). The processing stage outputs two types of CSV files: time series of signal intensities from each recorded channel and ROI length. CSV files enter the analysis stage (blue) to generate normalized fluorescent traces and detect peaks along with other signal processing metrics. These initial traces can be visualized to curate the data. Curation generates a configuration file that works as metadata across platforms and users. Curated data can be reanalyzed and used to visualize final data and compare across groups (yellow). Analysis and output stages are performed in the GUI (red dashed box), along with other metrics. Dashed arrows indicate optional steps.

The first pipeline step converts the raw data to TIF format, thereby avoiding compatibility issues that may arise when parsing different proprietary formats (Figure 1). All embryos are then segmented using histogram equalization, followed by intensity threshold binarization (Otsu 1979). Boxes surrounding the segments are cropped into individual time-lapses for each embryo. Cropping results in a substantial memory reduction, as most background pixels are removed, with cropped images typically accounting for around 40% of the original size.

The next step is to process each individual specimen. First, the ROI, which in our case is the entire central nervous system (CNS), is defined by binarizing the structural channel and selecting the largest connected component. This process is repeated at every time point to generate an “adaptive ROI”. From these adaptive ROIs, the average signal intensity for both channels is extracted. The results are saved as CSV files and are the basis for downstream analysis.

The ROI is also used to measure the length of the CNS (Figure 2). Drosophila embryo CNS length serves as an internal proxy for neurodevelopmental stages, enabling more accurate comparisons across embryos (Carreira-Rosario et al. 2021). The CNS length is calculated by centerline estimation. First, a distance transform is applied to the binarized image, and local maxima points are detected. Depending on the embryo’s orientation, some points may be part of the brain lobes and must be filtered out to accurately measure the CNS length. To obtain a robust centerline estimate that can ignore outliers, we use RANSAC (Fischler and Bolles 1981) over the local maxima points and measure the overlap between the fitted line and the binary image. CNS length is also detected frame by frame and exported as a CSV file (Figure 1).

**Figure 2:**
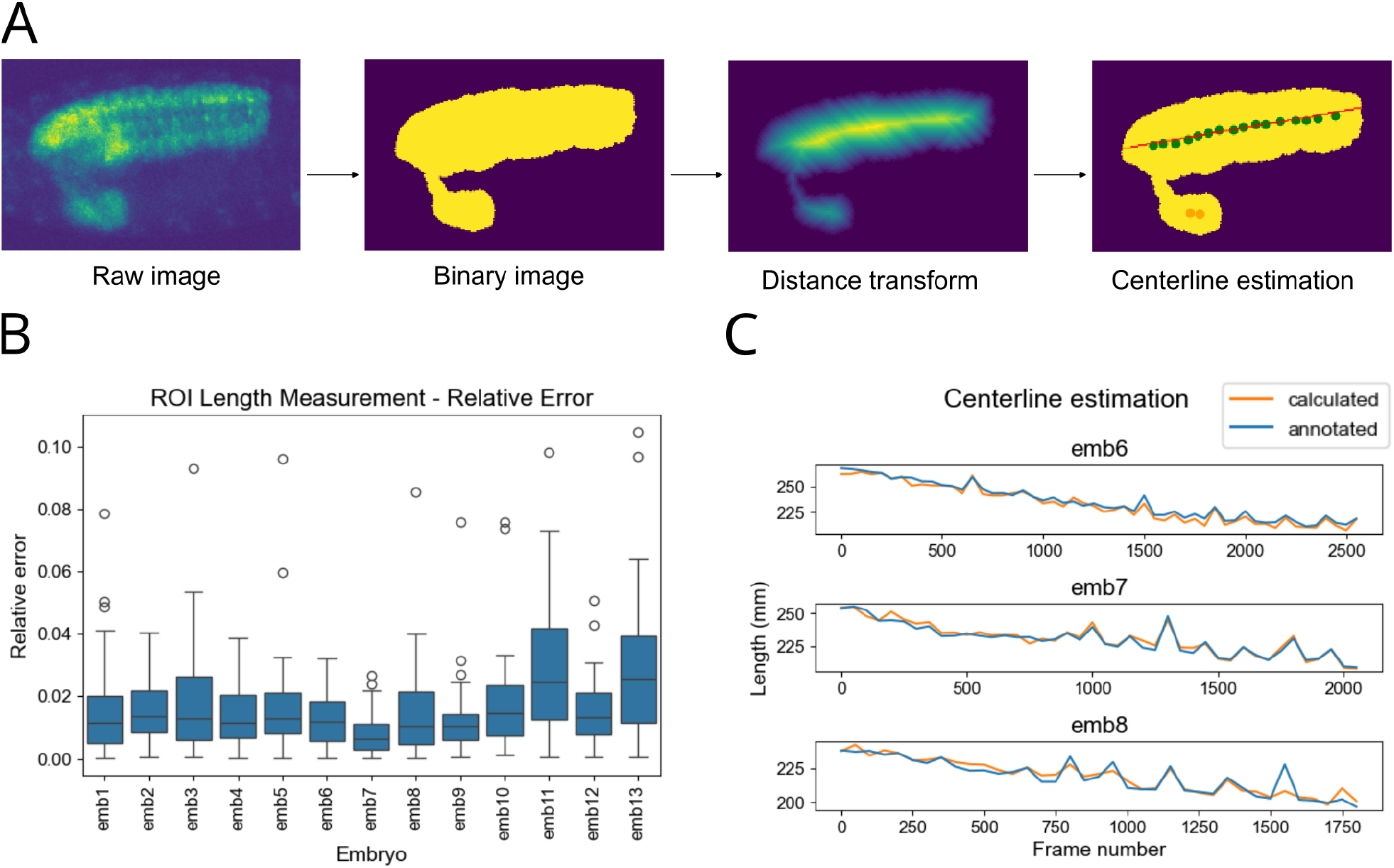
ROI length measurement algorithm and validation. A) Steps to calculate the ROI length. The ROI length is calculated by centerline estimation (red line) using local maxima (dots) in the distance transform, followed by RANSAC to ignore outliers (orange dots). B) Validation of the method as relative error (measured - annotated) / annotated. Each whisker bar summarizes the relative error for frames taken at intervals of 50 timepoints. C) Absolute values over a time series for three representative embryos.

The package utilizes average signal intensity measurements to calculate ΔF/F traces and peaks. For ΔF/F, we first calculate the ratiometric signal (dynamic signal / structural signal) and then its baseline, which is defined as the average of the N lowest values within a sliding window. The generated ΔF/F traces contain long-duration bouts of activity with superimposed fast transients (Figure 3). The former represents the bursts of activity and is the most relevant for the initial analysis. To mark only these more prolonged bouts, we apply a low-pass frequency filter to omit transients. Peaks in the filtered trace are detected using SciPy (Virtanen et al. 2020). Finally, the detected peaks are ported to the original ΔF/F signal.

**Figure 3:**
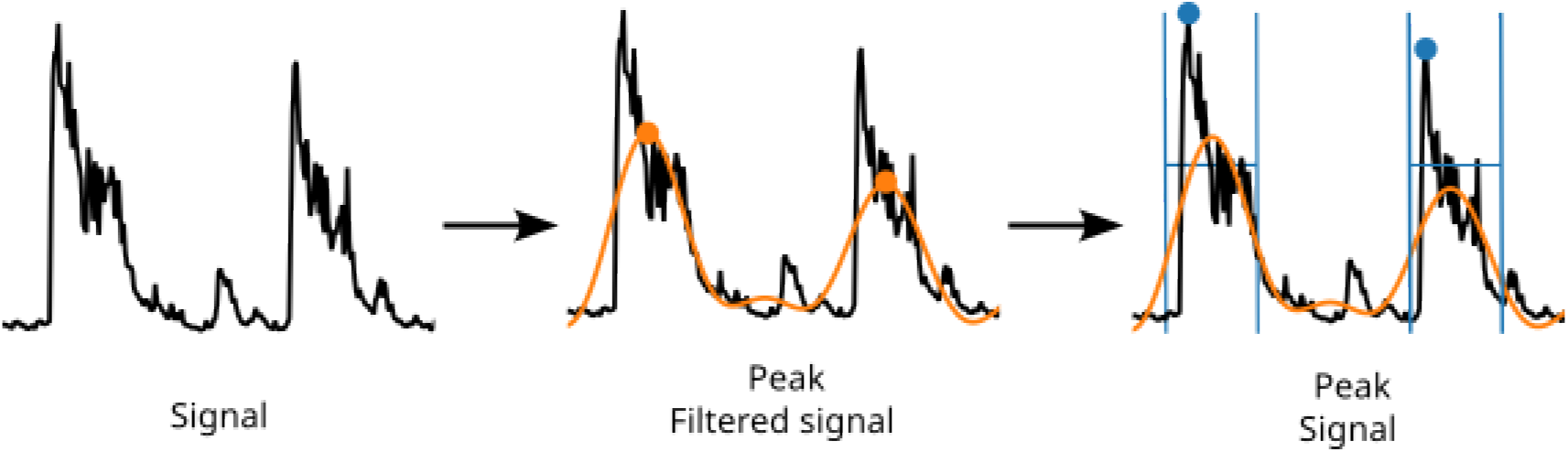
Peak detection algorithm. A low-pass filter (orange line) is applied to the ΔF/F signal (black line) to remove fast transients. The peak in the filtered signal (orange dot) is then ported back to the ΔF/F (blue dot) signal by selecting the leftmost peak within a search window (blue lines).

Results can be visualized and curated in a GUI implemented in `PyQt6` (Figure 4). During curation, researchers can modify data analysis parameters, which are persisted in a JSON configuration file and utilized by the core analysis code across different machines and users. Finally, a large number of different metrics and representations derived from ΔF/F, CNS length, and peaks can be visualized and plotted using the GUI. These include SNA onset, burst duration and spectrograms, among others.

**Figure 4:**
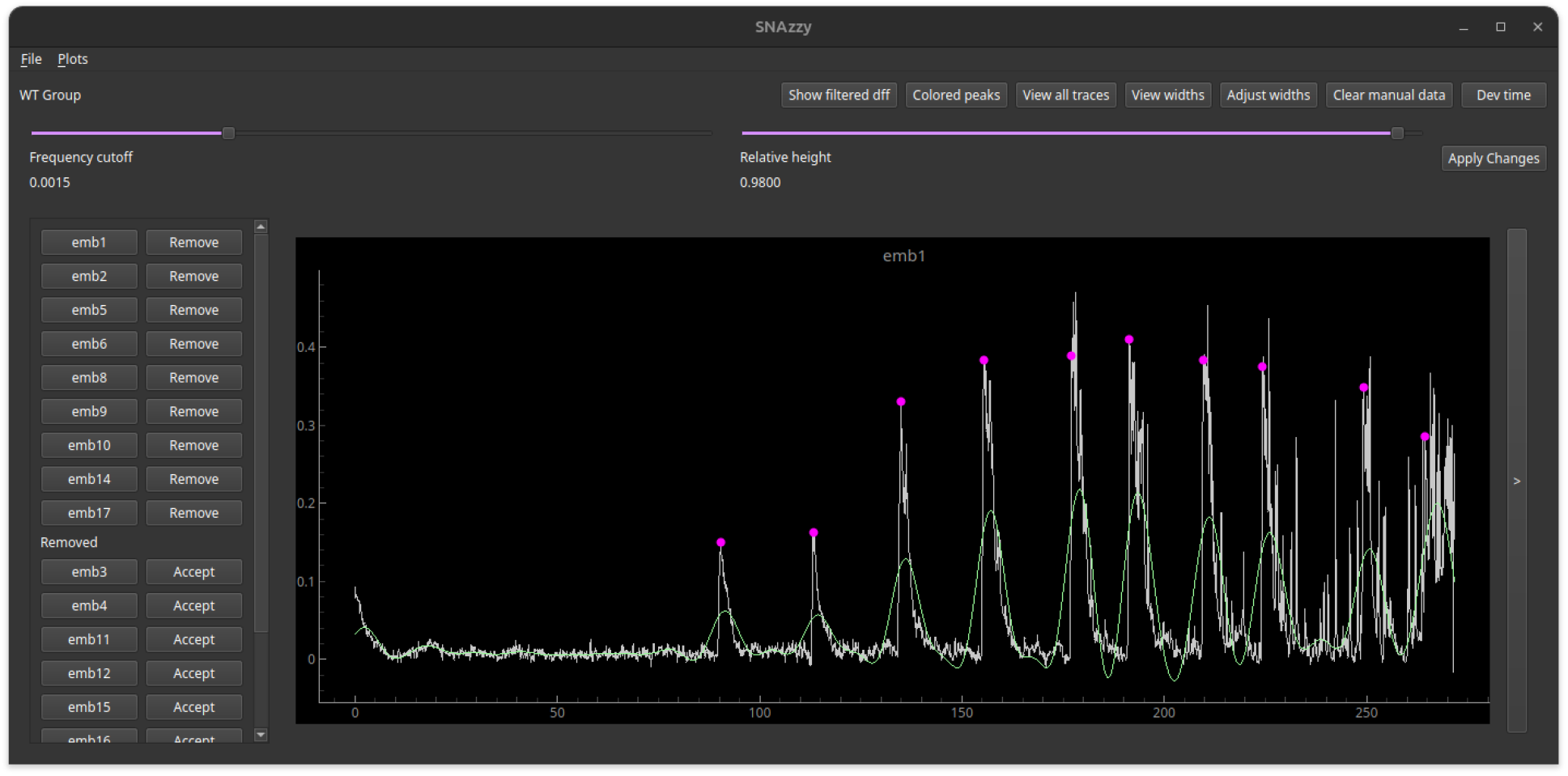
GUI for data validation, curation, visualization and plotting. Initial GUI screen. A ΔF/F trace (white) and the corresponding peaks (magenta dots) are shown. The low-pass filtered signal (green line) is used as a reference to determine peaks. The GUI enables the modification of analysis parameters, visualization of data, and comparison of metrics across groups of experiments, as well as manual adjustment of peak data.

## Conclusion

In conclusion, genetically encoded fluorescent indicators and microscopy systems are evolving rapidly, increasing the data acquisition throughput. Custom open-source tools are needed to handle such large data files. `SNAzzy` addresses this by offering an automated, scalable, and user-friendly platform for analyzing synchronous network activity in developing embryos. As an open and versatile solution, `SNAzzy` offers tools for a broader range of applications in time-lapse fluorescence imaging across diverse biological systems.

## AI Usage Disclosure

No generative AI tools were used for the development of the software. GPT 6 Astra was used to refine the documentation and minor grammar corrections in the manuscript. The authors have reviewed and validated all suggested changes and bear full responsibility for the final work.

## Acknowledgments

We acknowledge Newton PenkoffLidbeck and D. Berfin Azizoglu for feedback on the manuscript. This work was partially funded by NINDS and the BRAIN initiative (R00NS119295) and NSF-GRFP (DGE2139754) (AJE).

## Notes

### Competing Interest Statement

The authors have declared no competing interest.

